# 3D point cloud of normalized difference vegetation index (NDVI) of segmented fruit and leaves in apple production

**DOI:** 10.1101/2022.10.24.513567

**Authors:** Nikos Tsoulias, Kowshik Kumar Saha, Manuela Zude-Sasse

## Abstract

A feasible method to analyse fruit at the plant considering its position, size, and maturity are requested in precise production management. The present study proposes the employment of light detection and ranging (LiDAR) to measure the position, quality-related size, and maturity-related chlorophyll of fruit.

During fuit development, apples were analysed in the laboratory (n = 270) with two LiDAR laser scanners measuring at 660 and 905 nm. From the two 3D point clouds, the normalized difference vegetation index (NDVI) was calculated. The correlation analysis with chemically analysed fruit chlorophyll content showed R^2^ = 0.81 and 0.02 % RMSE.

The method was validated on 3D point clouds of 12 fruit trees in the orchard. Segmentation of individual apples was carried out during fruit development on five measuring dates, validated with manual rating (n = 4632). The non-invasively obtained field data showed good calibration performance capturing fruit position, fruit size, fruit NDVI of R^2^ = 0.99, R^2^ = 0.97, R^2^ = 0.71, respectively, considering the related reference data.

For 3D data of leaves, earlier shown analysis of leaf area and leaf chlorophyll by means of LiDAR was confirmed. The new approach of non-invasive laser scanning provided physiologically and agronomically valuable time series data on differences in fruit chlorophyll affected by the leaf area to fruit ratio, as well as differences of fruit chlorophyll in different growing position at the tree. Resulting, the method provides a tool for production management, e.g. crop load management, and integration in harvest robots.

## Introduction

Consumption of fresh fruit is recommended and worldwide an estimate of 883 million t, with a share of 10 % for apple, were produced in 2020 (FAO, 2021). Consumption of apples (*Malus* x *domestica* Borkh.) is globally high regardless of local production capability resulting in apple being one of the most traded fruit in the world. During ontogenesis of apple, the harvest maturity is determined by the complex ripening process of the fruit, which affects the fruit storability and market quality (Jones et al., 1965; OECD, 2018). Apples harvested too early may cause reduced blush colour, lack of cultivar-specific aroma, and harsh cortex tissue, whereas too late harvest reduces storability resulting in postharvest loss and food waste along the supply chain. Apple is a climacteric fruit, showing enhanced ethylene production and respiration rate at commercial harvest (Nelson, 1940; Biale, 1964; Hewitt and Dhingra, 2020). In the climacteric phase, the chlorophyll content decreases and, therefore, the change of chlorophyll during fruit development can be employed as an indicator for maturation and ripening (Knee, 1980; Zude-Sasse et al., 2002; Han et al., 2018). Visual monitoring of apple peel colour is widely performed to identify the ripening stage at harvest. The instrumental, non-destructive analysis of total fruit chlorophyll is established based on colorimetry as employed for inline grading and analysis of fruit homogeneity considering its colour appearance (Walsh et al., 2020a). Here, the chlorophyll content is represented in the red channel of red, green, blue (RGB) cameras or, e.g., along the a* axis of L*a*b* color space. Spectrophotometry for fruit pigment analysis (Knee, 1980; Merzlyak et al., 1999; Zude-Sasse et al., 2002) became commercially available as handheld systems (Walsh et al., 2020a) and for inline grading to obtain information on the fruit maturation. Furthermore, spectraloptical analysis with enhanced wavelength resolution enables the measurement of individual chlorophylls absorbing in the red wavelength range, such as chlorophyll_a, _b, and pheophytin_a (Seifert and Zude-Sasse, 2016). Such non-destructive methods can be applied in laboratory or packhouse conditions, however results are affected by varying lighting conditions appearing in field applications.

In outdoor measurements, the normalized difference vegetation index (NDVI) was introduced using the sun as light source and measuring the reflectance at Q band of chlorophyll absorption (Rouse et al., 1973). The NDVI examines the difference/sum ratio of reflectance in the red wavelength range, frequently at 660 nm, and shortwave near infrared (NIR) radiation bands varying between 700 nm and 900 nm (NIR-RED)/(NIR+RED). This index was initially used to assess the presence of vegetation and its application was confirmed in many studies (Zhou et al., 2001; Anyamba and Tucker, 2005; Tucker et al., 2005). The NDVI and other vegetation indices were employed in remote sensing approaches such as satellite imaging for estimating the vegetation cover, leaf area index (Bannari et al., 2009), and chlorophyll content of tree canopy (Li et al., 2018). Other studies have considered multispectral cameras mounted on unmanned aerial platforms to relate the canopy NDVI with tree vigor (Ballester et al., 2018; Ampatzidis et al., 2019), yield and quality prediction (Kasimati et al., 2022) of fruit trees.

The NDVI of fruit was introduced and compared to whole spectrum analysis for predicting the fruit chlorophyll content considering the fruit surface area (Zude, 2003). Again, various indices were tested (Zude-Sasse et al., 2002; Ziosi et al., 2008). More advanced, the use of time-resolved spectral analysis enabled to separate the absorption and scattering effects of the fruit tissue and, by working on the absorption coefficients, enhancing the accuracy of NDVI analysis, particularly, in stone fruit (Seifert et al., 2015). In apple, the NDVI measured in remittance geometry with contact to the fruit was reported as robust for predicting the fruit chlorophyll (Kuckenberg et al., 2008; Seifert et al., 2016). However, all methods are requested to be carried out in close proximity to the fruit and varying light conditions alter the signal. The potential usages of the chlorophyll sensor on a robot for crop load management and harvest request the classification of various chlorophyll classes on the tree capturing the distance between the row, where the robot is placed, and the fruit in the canopy, which may be assumed as 0.3 – 1.6 m.

Three-dimensional (3D) vision systems, such as light detection and ranging (LiDAR) laser scanning may overcome the limitations of established point or 2D imaging methods (Gongal et al., 2018; Walsh et al., 2020b; Keller et al., 2022). So far, LiDAR is used in remote sensing application in arable farming and forestry due to its capability to provide 3-dimensional (3D) geometric information of vegetation in field conditions (Deery et al., 2021; Zhu et al., 2021). Generally, a monochromatic laser emitting in the visible or short wave near infrared wavelength range, is employed as light source of LiDAR systems. The receiver acquires distance information from sensor to object by means of triangulation, which creates a set of points when the beam hits the surface of an object in space leading to 3D point clouds. Advancement of terrestrial LiDAR sensors facilitate to acquire also the intensity of backscattered reflection at each point measured. Thus, besides geometric information, intensity of reflected signal becomes available as shown earlier for the segmentation of apple fruit (Tsoulias et al., 2020). Furthermore, estimation methods for structural parameters and chlorophyll content in broadleaf plants were developed employing terrestrial LiDAR data (Eitel et al., 2010). Subsequently, simulation studies were published to quantify chlorophyll content of broadleaf forest trees (Cifuentes et al., 2018). Several studies proved the application of LiDAR 3D point cloud with intensity information for detecting the overall chlorophyll status of vegetation (Watt and Donoghue, 2005, Clawges et al., 2007, Thoren and Schmidhalter, 2009). Despite the physiological importance of fruit chlorophyll and the known relationship of fruit chlorophyll content with the NDVI, the NDVI of segmented fruit and its development during the growth season weren’t reported based on LiDAR data so far.

Therefore, objectives of the present study were to (i) develop a method for estimating the segmented NDVI of foliage and fruit obtained with LiDAR at 660 and 905 nm (NDVI_LiDAR_), (ii) verify the relation of temporally measured NDVI_LiDAR_ with the fruit chlorophyll content (iii) characterizing the relationship of fruit NDVI_LiDAR_ and leaf area to fruit ratio and on leaf area to fresh mass ratio.

## Materials and methods

### Site description

The experiment was conducted in the Leibniz Institute for Agricultural Engineering and Bioeconomy (ATB) experimental apple orchard located in Potsdam-Marquardt, Germany (Latitude: 52.466274° N, Longitude: 12.57291° E). The field is located on an 8 % slope with southeast orientation, planted with trees of *Malus* × *domestica* Borkh. ‘Gala-Brookfield’ and JonaPrince’, and pollinator trees ‘Red Idared’ each on M9 rootstock with 0.95 m distance between trees, trained as slender spindle with an average tree height of 2.8 m. Trees are supported by horizontally parallel wires.

All measurements were conducted five times during fruit development. Trees were scanned 65 days after full bloom (DAFB_65_) during the end of cell division stage of fruit, before the Brookfield-typical red blush colour (Sadar, Urbanek-Krajnc, & Unuk, 2013) appeared, subsequently during fruit development until commercial harvest maturity, 75, 90, 105, and 130 days after full bloom (DAFB_75_, DAFB_90_, DAFB_105_, DAFB_130_, respectively). Twelve trees of Gala-Brookfield were analysed by means of the non-invasive sensor system and fruit were sampled from these trees for the lab measurements. Further neighboring trees were used for destructive measurements such as leaf area analysis after defoliation.

### LiDAR measurements in field and laboratory conditions

#### Field data acquisition

A phenotype sensing system was mounted on a circular conveyor platform, established in the experimental apple orchard (TechGarden, ATB), employing an electrical engine working with 50 Hz (DRN71, SEW Eurodrive, Germany) and stainless-steel chain with mechanical suspensions for varying plant sensors (Figure 1a). Two mobile 2D LiDAR sensors emitting at the wavelength of 905 nm (LMS-511, Sick AG, Waldkirch, Germany) and of 660 nm (R2000, Pepperl Fuchs, Mannheim, Germany) were mounted vertically on the metal frame at 0.7 m above the ground level (Figure 1b). The LMS-511 and R2000 were configured with 0.1667° and 0.029° angular resolution, 25 and 20 Hz scanning frequency, scanning angle of 180° and 270°, respectively. A real time kinematic global navigation satellite system (RTK-GNSS; AgGPS 542, Trimble, Sunnyvale, CA, USA) was used to geo-reference the data and an inertial measurement unit (IMU; MTi-G-710, XSENS, Enschede, Netherlands) to acquire orientation information were placed on the sensor frame. The IMU was placed 0.3 m aside from the LiDAR sensor, while the receiver antenna of RTK-GNSS was mounted 0.6 m above the laser scanner. The platform enables the automated monitoring of 109 trees in one row of 84 m length, of which 12 trees were analysed in this study considering field and lab data.

**Figure 1:**
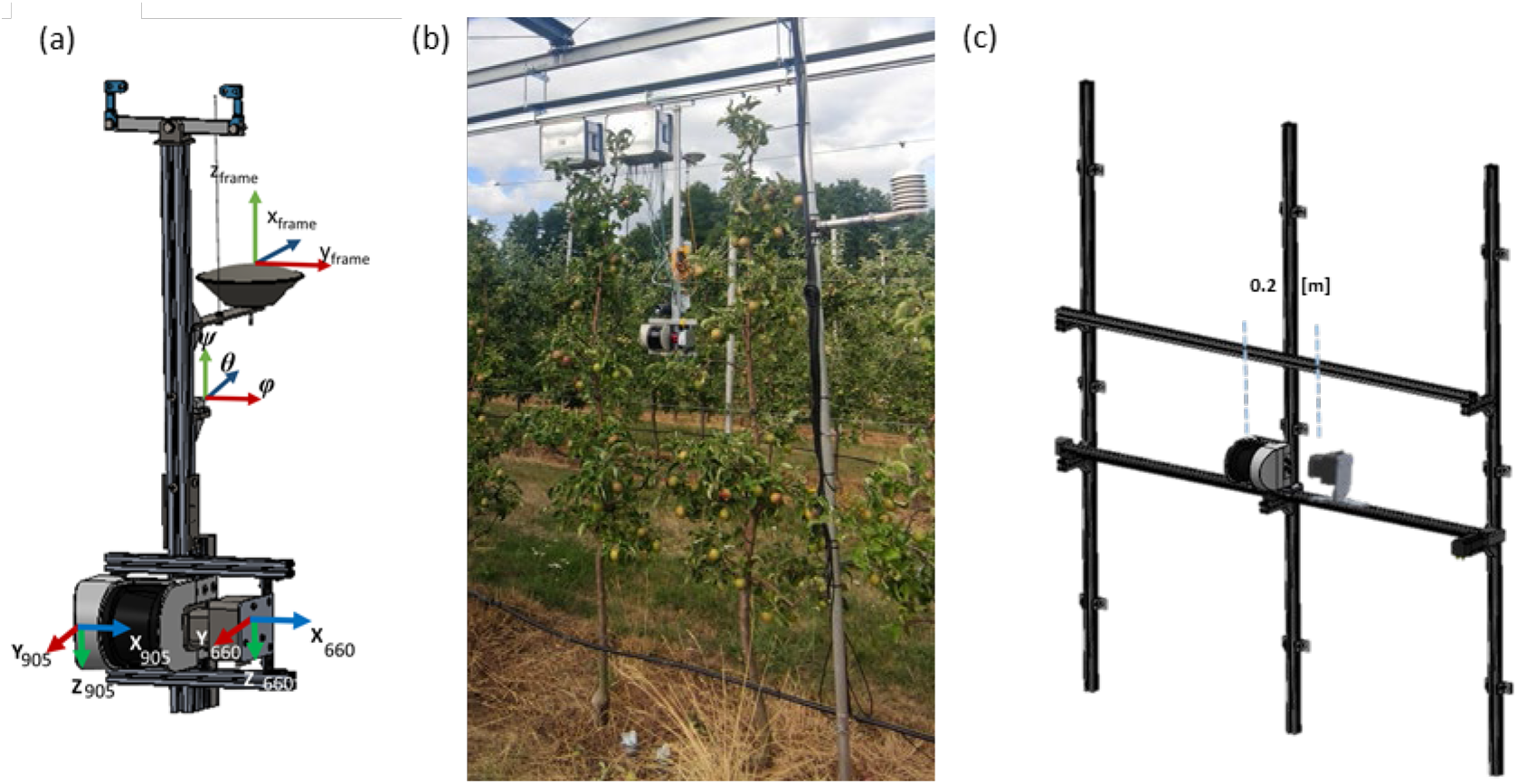
Sensor frame (a), mounted on chain conveyor in the experimental orchard (b), mounted on a tooth-belt conveyor in dark room.

#### LiDAR laboratory data acquisition

After each measurement with the phenotyping sensor system, apple samples (n = 45) were collected. A subsample of apples (n = 10), from the 45 apple-batch, was additionally scanned in the laboratory on the five measuring dates. The measurement was carried out in a dark room, controlled ventilation and temperature. A rigid linear toothbelt conveyor system (Module 115/42, IEF Werner, Germany) of 2 m length, employing a servo positioning controller (LV-servoTEC S2, IEF Werner, Germany), was mounted on a rigid aluminium frame to carry the two LiDARs and scan each apple individually (Figure 1 c). The linear conveyor moved at 20 mm s^-1^ (± 0.05 mm accuracy) forward speed.

### LiDAR data processing

#### Point cloud reconstruction

The 3D point cloud dataset was generated and processed in the Computer Vision Toolbox™ of MATLAB (2018b, Mathworks, Natick, MA, USA). Rigid translations and rotations were applied on each point of 3D cloud, while alignment of pairing tree sides was carried out with iterative closest point algorithm (Tsoulias et al., 2019). Trees were segmented based on stem position and planting distance to gain points per tree (PPT) from each plant (Tsoulias et al., 2019).

In the laboratory, data were recorded and all measured distances were filtered to remove surrounding points. As the apple samples were scanned from 0.9 m distance, the distance filter was configured between 0.75–1.25 m. Distant filtering also helped to reduce the raw data file size and resulted in less processing time in further step. Using the corresponding distances in *x* and *y* direction in the vertical *x-y* plane considering the sensor captured distance. The linear movement of the LiDAR scanner was in *z* direction and displacement in this direction was calculated by forward speed and time difference between each vertical line of scan.

For all data sets from field and laboratory, the 3D point clouds at 660 nm and 905 nm were further processed capturing data sets of position in local Cartesian x, y, z coordinate system and reflected intensity of each point (R_ToF_).

#### Leaf area estimation

For each point of 3D tree cloud the geometric feature of linearity (L) and curvature (C) were calculated applying the k-nearest neighbours (KNN) algorithm in the local neighbourhood of points P_i_= [x_i_; y_i_; z_i_] (Tsoulias et al., 2022). The total number of P_i_ within each tree’s cloud was used to estimate the mean of all nearest neighbors 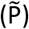. The latter was used to produce the eigenvalues (λ_1_, λ_2_, λ_3_), after the decomposition of covariance matrix. The probability density function was performed to define the thresholds of linearity, curvature and LiDAR’s backscattered intensity (R_ToF_) to distinguish the 3D points of woody parts (W) from leaves. The value with the highest likelihood (mode) within the R_W_, C_W_, and L_W_ classes was used as threshold (*R_th,W_; C_th,W_*; and *L_th,W_*). Points which fulfilled the criteria of L_W_ ≤ *L_th,W_, C_th,W_* ≤ C_W_, and *R_th,W_* ≤ *R_ToF,W_* were segmented and categorized as wood.

Segmented points of wood were subtracted from the total number of PPT. A linear regression model was built to express the relationship between manually obtained leaf area data and remaining PPT, separately for each growth stage. The model performance was evaluated based on the adjusted coefficient of determination in calibration (R^2^_adj_) and and root mean squared error (RMSE). The linear calibration was applied to convert PPT into LA_LiDAR_ of each tree (Tsoulias et al., 2022).

#### Apple detection, counting, and sizing

For defining the position and shape of apples, the geometric feature of curvature (C) and R_ToF_ were used considering each point of the 3D tree point cloud (Tsoulias et al., 2020). Following the previous method, the local neighbors were decomposed and eigenvalues were produced. The values closer to 100, the higher the likelihood for shape of point appearance to be curved. Threshold values of apple points in terms of C and reflected intensity (C_A_ and R_ToF,A_) were defined according to Tsoulias et al., (2020) by performing probability density function. The points that fulfilled the criteria of C_th,A_ ≤ C_A_, and R_th,A_ ≤ R_ToF,A_ were segmented and categorised as apples. Subsequently, a density-based scan algorithm (DBSCAN) was applied to find the point sets, using the mean manually measured diameter of apples that was found in the neighborhood search radius (*ε*) and the value 10 as a minimum number of neighbors. The value of 10 was applied resulting from manually run tests showing that less neighboring points result in random appearance of sets. The maximum distance in x and y axes of fruit points was considered as the diameter of each point set recognized as an apple (D_LiDAR_). Thereafter, *k*-means clustering was applied to find the fruit center(s) in each apple cluster and count the number of fruit per tree (Fruit_LiDAR_).

Subsequently, the ratio of leaf area to number of fruit per tree (LA_LiDAR_:Fruit_LiDAR_) was estimated in each growing stage in low and high sections of tree canopy determined by the wire structure at 1.8 m statically supporting the tree.

#### NDVI estimation

The point clouds of segmented apples was obtained from the two LiDAR laser scanners, which varied considering the point density due to different scanning frequency and angle resolution. The corresponding point clouds of the same apple were merged by means of a density histogram applied on segmented apple clouds (Figure 2).

**Figure 2:**
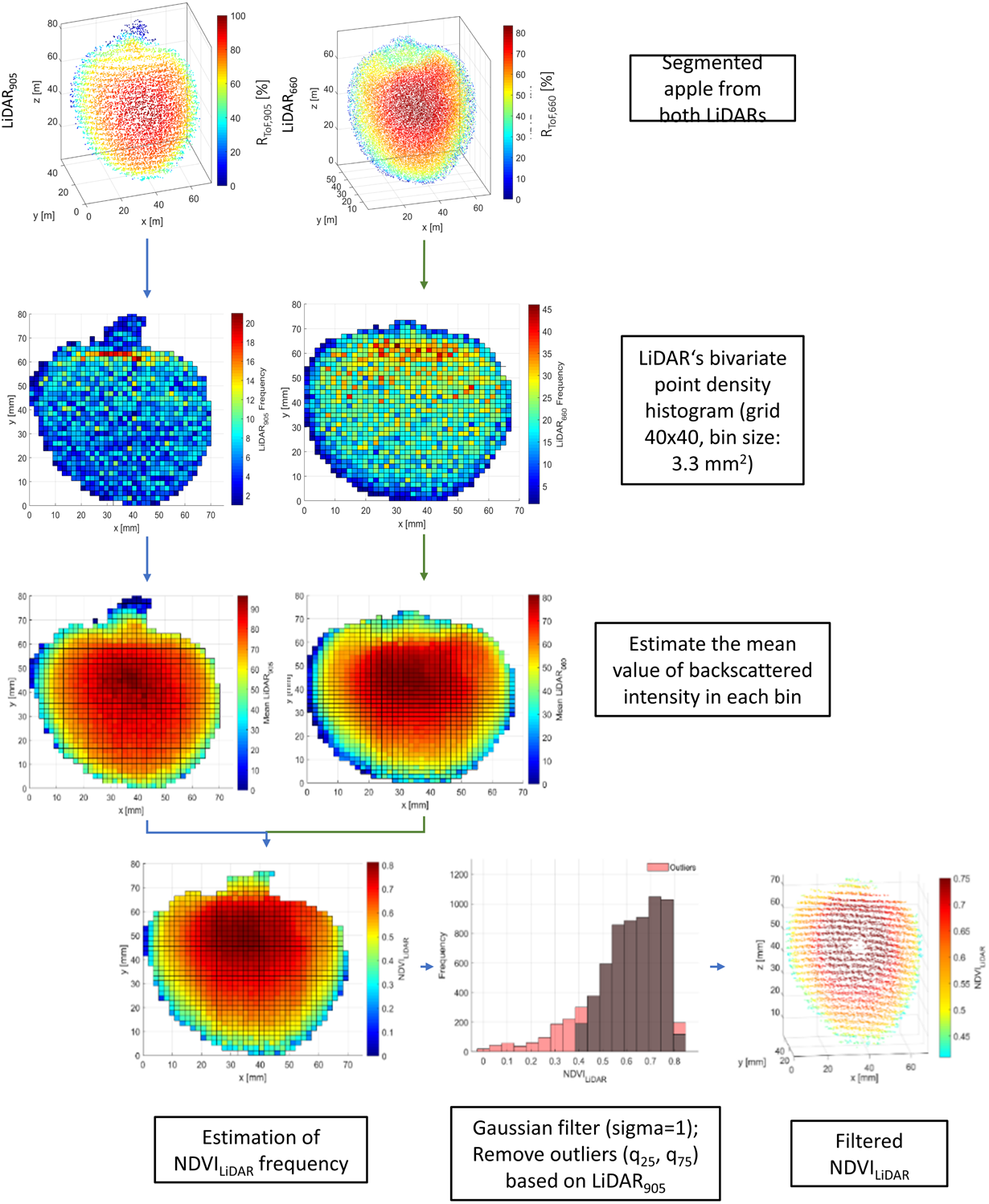
Flow chart to calculate the fruit NDVI_LiDAR_ from the segmented fruit point clouds.

The x and y values of each point paired in bins of size 3.3 mm^2^, allowing to describe the underlying point density distribution of apple R_ToF_ as means of each bin within the grid of 40 x 40 bins within the shape of each apple. The number of points in each bin varied according to the shape and size of apple. The bin’s mean values allowed to use the corresponding values of both LiDAR systems for calculating the normalised differential vegetation index (NDVI_LiDAR_).

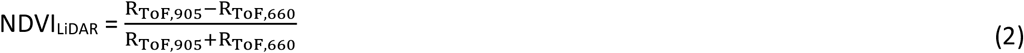

Where R_ToF,905_ and R_ToF,660_ represent the reflected intensity measured at 905 and 660 nm, respectively. The derived NDVI_LiDAR_ cloud of each apple was corrected for outliers by means of Gaussian filter with standard deviation of distribution equal to 1. The interquartile range, estimated by subtracting the points of lowest and highest quartile, were used in further calculations. The process was applied in all segmented fruit point clouds over the growing stages (Figure 2). The mean NDVI_LiDAR_ of each fruit was processed further. The results were categorised in low (.0 −1.8 m) and high (1.8-2.6 m) sections of tree heights determined by the wire structure.

### Reference measurements

#### Leaf area

On each measuring date, 3 apple trees neighboring the monitored 12 trees were manually defoliated. The overall 18 defoliated trees were scanned before and after defoliation. The obtained leaf area per tree was manually recorded (LA_Manual_) with a desktop scanner (Scanjet 4850, HP, USA) by counting the number of green pixels per leaf with a MATLAB script (Vers. 2018b, Mathworks, Natick, MA, USA). The area of 6241 pixel corresponded to 1 cm^2^. The results were categorised in low (.0-1.8 m) and high (1.8-2.6 m) section of tree heights determined by the wire structure.

#### Crop load and fruit quality

Fruit number was manually counted per tree (n = 12) in low and high sections of the canopy separated by the construction wires at 1.8 m on the last three measuring dates.

At the five measuring dates 45 fruit were analysed in the laboratory. Fruit diameter (D) [mm] was manually measured in the laboratory by means of a digital calliper gauge considering the mean diameter of two measurements taken equatorially with 90° difference. The fresh mass (FM) [kg] was measured by weighting each fruit sample. Moreover, measurements of diameter (n = 180) were recorded at the two tree height ranges, low and high over the growing stages.

The NDVI was recorded with a handheld spectrophotometer (Pigment Analyzer, PA1101, CP, Golm, Germany) in remittance geometry by placing the sensor probe directly on the fruit surface avoiding stray light with the silicon probe of the device. Also, these analyses were separated in low and high sections of tree canopy at 1.8 m.

The chlorophyll content of skin and associated hypodermis tissue (2 mm thickness) of apples (n = 45) was destructively analysed at each measuring date over the fruit development. The chlorophyll a and b and pheophytin a contents were analysed by spectrophotometry after acetone/diethyl ether extraction by means of iterative multiple linear regression considering the standard spectra of the three chlorophylls (Pflanz and Zude, 2008).

### Data evaluation

Descriptive statistics were applied to all datasets capturing minimum (min), maximum (max), mean, standard deviation (SD). A regression analysis was performed to quantify linear and logarithmic relationships between the manual measurements and LiDAR-derived data over the growing stages, and RMSE, mean bias error (MBE) were caluculated. Descriptive statistics were carried out by Matlab (v.R2018b, Mathworks Inc., Natick, MA, USA).

## Results and discussion

### Fruit segmentation

Slender spindle form the major training system of apple trees in world-wide production, providing a 3D structure in which the fruit are more or less evenly distributed according to the success of the thinning measure. Yield monitoring is considered as an important step when implementing precision horticulture techniques in orchards. In the present study, fruit detection technique was applied to extract the number of fruit from the the point cloud of LiDAR_905_ and LiDAR_660_ (Figure 3). The fruit segmentation routine was described earlier (Tsoulias et al., 2020) and marginal difference of noninvasively detected and manually counted fruit was found for both LiDAR systems (Figure 3).

**Figure 3:**
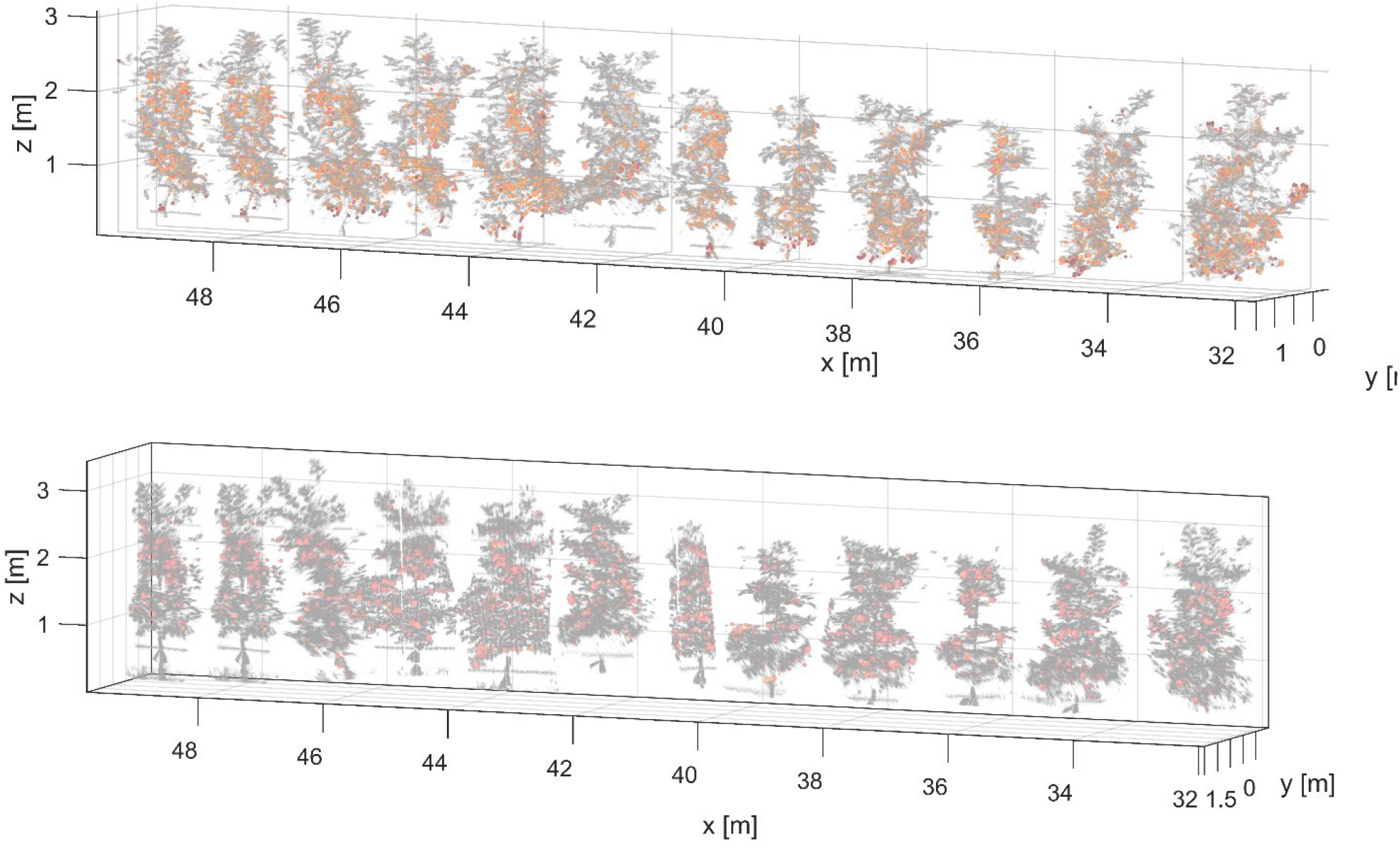
3D point clouds of 12 apple trees with the segmented fruit labelled in red, measured on the last measuring date at harvest. Measurements were done with LiDAR_905_ (above) and LiDAR_660_ (below).

The LiDAR_905_ was used as a pilot sensor to segment the fruit, pointing to R^2^ = 0.99 for the fruit detection (Table 1). The highest difference between manually and estimated number of fruit was observed at DAFB_65_, when mean apple size was D_Manual_ = 35.5 mm, reaching R^2^ = 0.88.

After the fruit localization, the estimated fruit size (D_LiDAR_) was compared to the manually measured diameter during the growth stages (Table 1). The detected diameter was related to the manual reading, especially in the latter stages, DAFB_105_ and DAFB_130_, when the fruit reached their maximum size resulting in R^2^ = 0.98 and 0.97 with RMSE = 0.34 % and 0.39 %, respectively. Generally, high measuring uncertainty was noticed on the first two measuring dates, when the fruit size was smaller. More specifically, a less pronounced relation was observed on the first measuring date, presenting an overestimation < 1 mm. Similar good results were reported earlier on the fruit detection and sizing at harvest (Gongal et al., 2018, Gene-Mola et al., 2019), whereas the early fruit sizing wasn’t approached frequently. Resulting, the LiDAR based analysis provides an accurate tool for early fruit counting.

**Table 1:** Reference (manual) data and LiDAR derived estimations with error analysis (mean bias error, MBE, root mean squared error, RMSE, coefficient of determination, R^2^) considering fruit number (Fruit_LiDAR_), fruit diameter (D_LiDAR_), and leaf area (LA_LiDAR_) measured at the tree in five growth stages of apples in day after full bloom (DAFB).

|  |  | DAFB<sub>65</sub> | DAFB<sub>75</sub> | DAFB<sub>90</sub> | DAFB<sub>105</sub> | DAFB<sub>130</sub> |
| --- | --- | --- | --- | --- | --- | --- |
| No.<br>$Fruit_{Manual}$ | Sum (n = 12, trees) | | | 795 | 771 | 771 |
|  | Mean | X | X | 65 | 64 | 65 |
|  | Max |  |  | 84 | 84 | 84 |
|  | Min |  |  | 19 | 19 | 19 |
| No.<br>$Fruit_{LiDAR}$ | Sum | 745 | 773 | 788 | 767 | 771 |
|  | Mean | 66 | 71 | 71 | 70 | 70 |
|  | MBE | -4 | -2 | -1 | 1 | 0 |
|  | RMSE (%) | 0.33 | 0.26 | 0.69 | 0.96 | 0.01 |
| | $R^2$ | 0.88 | 0.96 | 0.96 | 0.99 | 0.99 |
| $D_{Manual}$<br>(mm) | Mean | 35.5 | 48.3 | 59.1 | 61.4 | 76.1 |
|  | Max | 41.7 | 52.5 | 65.0 | 66.0 | 83.0 |
|  | Min | 32.0 | 41.4 | 52.0 | 56.0 | 68.0 |
| $D_{LiDAR}$<br>(mm) | Mean | 35.8 | 48.4 | 59.1 | 61.5 | 76.1 |
|  | MBE | 0.37 | 0.46 | 0.11 | 0.05 | -0.08 |
|  | RMSE (%) | 6.13 | 6.04 | 6.22 | 3.08 | 3.02 |
| | $R^2$ | 0.85 | 0.87 | 0.86 | 0.95 | 0.98 |
| $LA_{Manual}$<br>(m<sup>2</sup>) | Mean | 6.40 | 6.41 | 6.47 | 6.48 | 6.54 |
|  | Max | 6.48 | 6.51 | 6.54 | 6.55 | 6.56 |
|  | Min | 6.26 | 6.26 | 6.20 | 6.41 | 6.47 |
| $LA_{LiDAR}$<br>(m<sup>2</sup>) | Mean | 6.41 | 6.51 | 6.54 | 6.50 | 6.60 |
|  | MBE | 0.17 | 0.22 | 0.23 | 0.23 | 0.25 |
|  | RMSE (%) | 3.14 | 3.03 | 3.12 | 2.42 | 6.36 |
| | $R^2$ | 0.89 | 0.95 | 0.96 | 0.95 | 0.86 |

When starting the LiDAR analysis, the foliage of decidious trees was almost developed. The mean of LA_LiDAR_ increased from 6.40 m^2^ to 6.54 m^2^ during the measuring period capturing the range between DAFB_65_ and DAFB_130_, respectively (Table 1). LA_LiDAR_ showed an overestimation in all growths stages, presenting a MBE of 0.17 m^2^ at DAFB_65_ and 0.25 m^2^ at harvest. The manually measured leaf area was correlated with LA_LiDAR_ over the measuring period, showing an R^2^ of 0.89 with RMSE = 0.03 % at DAFB_65_ and R^2^ = 0.86 with RMSE = 0.06 % at DAFB_130_. The accuracy of the non-invasive analysis was limited due to occlusions and coinciding leaf surfaces as suggested in many plant species (Deery et al., 2021; Keller et al., 2022). However, such data derived from LiDAR point clouds are informative for late hand thinning considering the leaf area to fruit ratio in crop load management (Penzel et al., 2021). According to the findings presented in Table 1, both information can be obtained with the same sensing technique.

### Bivariate point density histogram at two wavelengths measured in laboratory and field conditions

The backscattered reflectance intensities measured with the two LiDAR sensors LiDAR_905_ (RToF,905) and LiDAR_660_ (R_ToF,660_) were extracted from apple point clouds (Figure 2) measured in the laboratory and in the orchard over the fruit growing stages. Applying the bivariate histogram of apple point clouds allowed to acquire the mean backscattered reflectance with the same point density from both laser scanners in the dark room and in the field (Figure 4). The number of points per apple increased with fruit size during the growth period, while the number of bin remained the same allowing a direct comparison between all fruit. R_ToF_ values found in previous work done on apple fruit, aimed at fruit segmentation, ranged between 60 % and 90 % (Gené-Mola et al., 2019) measured in the shortwave near infrared range of 905 nm. The same range was found in the present study for R_ToF,905_ (Figure 4). Comparing lab and field data, a sharper peak between 60 % and 80 % was found in field measurements. In field conditions, apple surface was captured by a lower number of points due to occlusions. The frequency distribution of R_ToF,905_ changed slightly over the fruit growth period.

**Figure 4:**
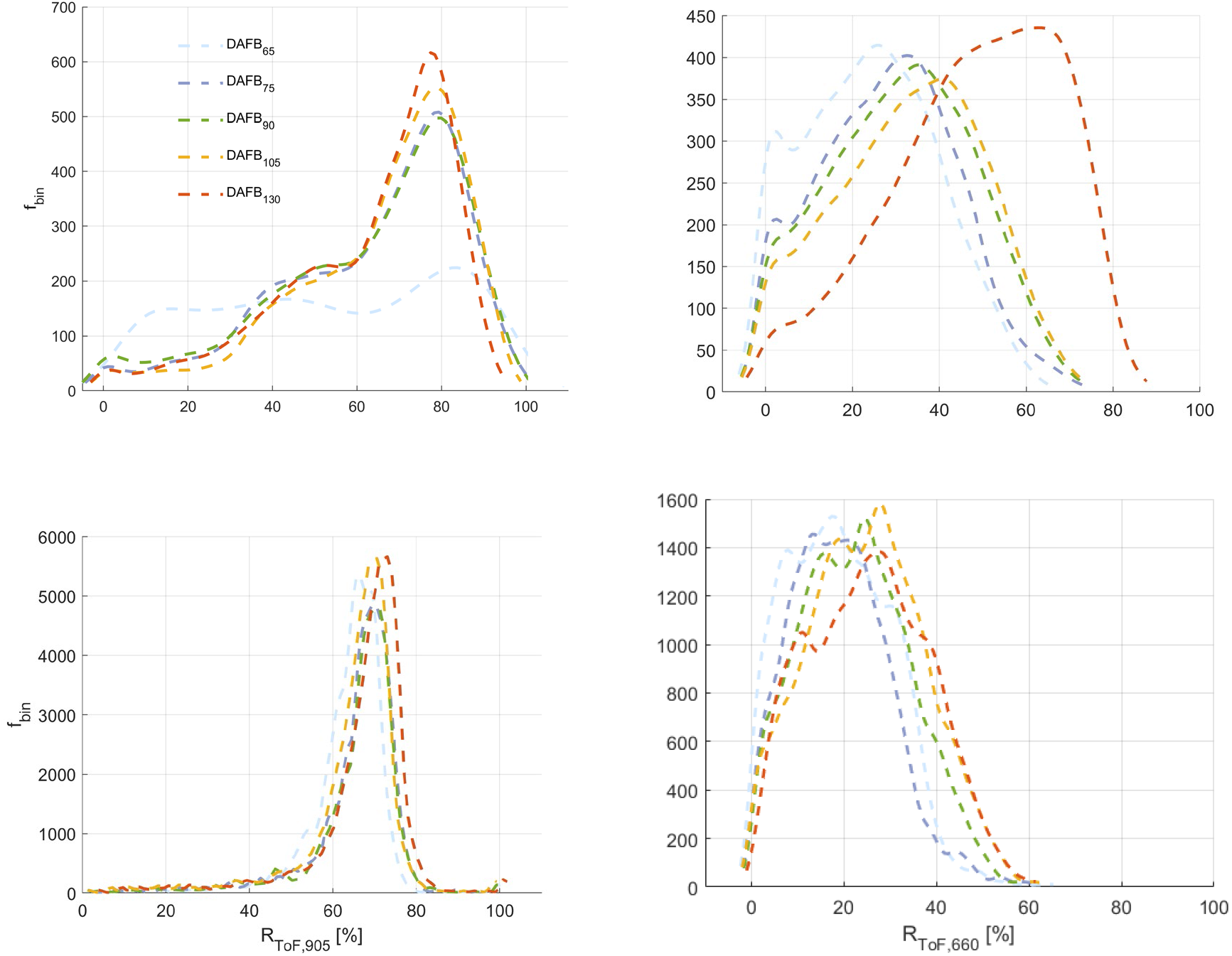
Bivariate histogramm considering all bins(n = 160 for each apple) of backscattered reflectance histogramm (R_ToF_) at five ripening stages measured in the lab (upper) (n = 10) and field (lower) (n = 771) for LiDAR_905_ (left column) and LiDAR_660_ (right column).

R_ToF,660_ ranged from 0 to 80 % in the lab, whereas in field conditions the narrowed range between 0 and 40 % was captured. The R_ToF,660_ showed lower values compared to R_ToF,905_ due to chlorophyll absorption (Figure 4). Furthermore, frequency curves were moving in direction from low to high intensity due to degradation of chlorophyll pigments which corresponds to reduced absorption coefficient in riper fruit. The most frequent value in R_ToF,660_ was found at 26.9 %, 33.8 %, 35.7, 41.3 %, and 63.6 % for DAFB_65_, DAFB_75_, DAFB_90_, DAFB_105_, and DAFB_130_, respectively. However, in field conditions high overlap of R_ToF,660_ intensity curves were found, with the most frequent value fluctuating with 18.1 %, 13.6 %, 23.5 %, 28.2 %, 27.9 % at DAFB_65_, DAFB_75_, DAFB_90_, DAFB_105_, DAFB_130_, respectively.

The frequency curves of resulting fruit NDVI_LiDAR_ show a high variance and, during the growth period, moved in the direction from low to high intensity as can be assumed due to degradation of chlorophyll pigments which corresponds to the fruit ripening process (Figure 5, left). More specifically, the NDVI_LiDAR_ ranged from 0.05 to 0.72 with a peak value of 0.34 at DAFB_65_. The NDVI_LiDAR_ values were reduced following chlorophyll degradation, reaching a peak value of 0.11 and a range between −0.25 and 0.38 at DAFB_130_. On the other hand, a reduced range was revealed over the growing stages in the NDVI_LiDAR_ measured in the orchard (Figure 5, right), again assumingly due to less points capture from the fruit in the partly overlapping foliage. During the first measuring days the fruit NDVI hardly changed. The last measuring date showed a clear decrease of NDVI_LiDAR_ in the laboratory as well as in field measurements. Such findings, obtained non-invasively, are consistent with previous findings measured with spectroscopy with the sensor probe being in contact to the fruit surface (Zude-Sasse et al., 2002; Muresan et al., 2017).

**Figure 5:**
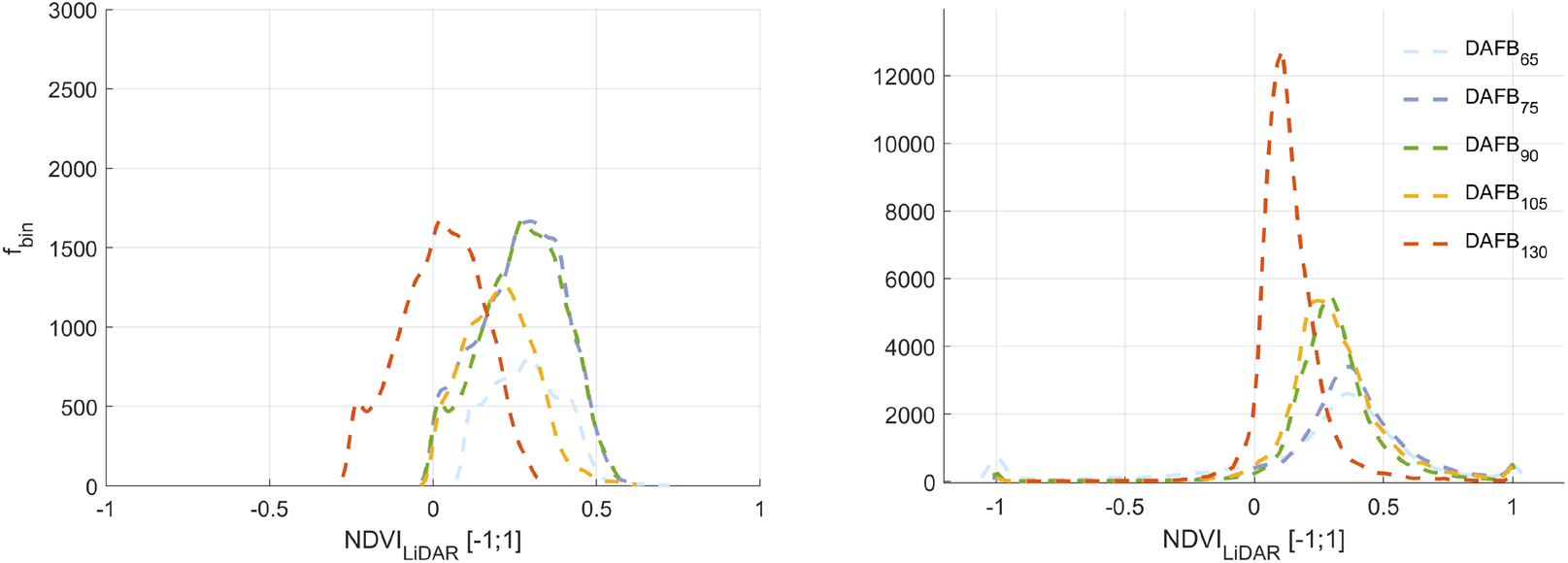
Bivariate histogramm of NDVI_LiDAR_ considering all bins (n = 160 for each fruit)] measured in dark room (left) and field (right) at five ripening stages.

### Correlation of NDVI_LiDAR_ and fruit reference analyses

For referencing, the NDVI_Ref_ was measured with a handheld spectrophotometer and the chlorophyll content (Chl_A_, Chl_B_, Chl_pheo_ and Chl_A+B+pheo_) were chemical analysed were recorded (n = 45 on each measuring date) over the growing stages after LiDAR scanning (Figure 6). The temporal decrease of NDVI was revealed, presenting a mean value of 0.88 with ± 0.05 standard deviation (SD) at DAFB_65_ and a mean value of −0.21 with a ± 1.21 SD at DAFB_130_. The variability of Chl_A_, Chl_B_ Chlpheo and its sum Chl_A+B+pheo_ were enhanced at DAFB_65_ and DAFB_75_, decreased among the following growth stages. Single factor ANOVA revealed clear difference (*p*< 0.001) among all ripening stages considering the spectrophotometrically measured NDVI_Ref_. The mean values for the total chlorophyll content (Chl_A+B+pheo_) were 0.98, 0.63, 0.46, 0.43 and 0.41 mg cm^-2^ at DAFB_65_, DAFB_75_, DAFB_90_, DAFB_105_, and DAFB_130_, respectively (Figure 6).

**Figure 6.**
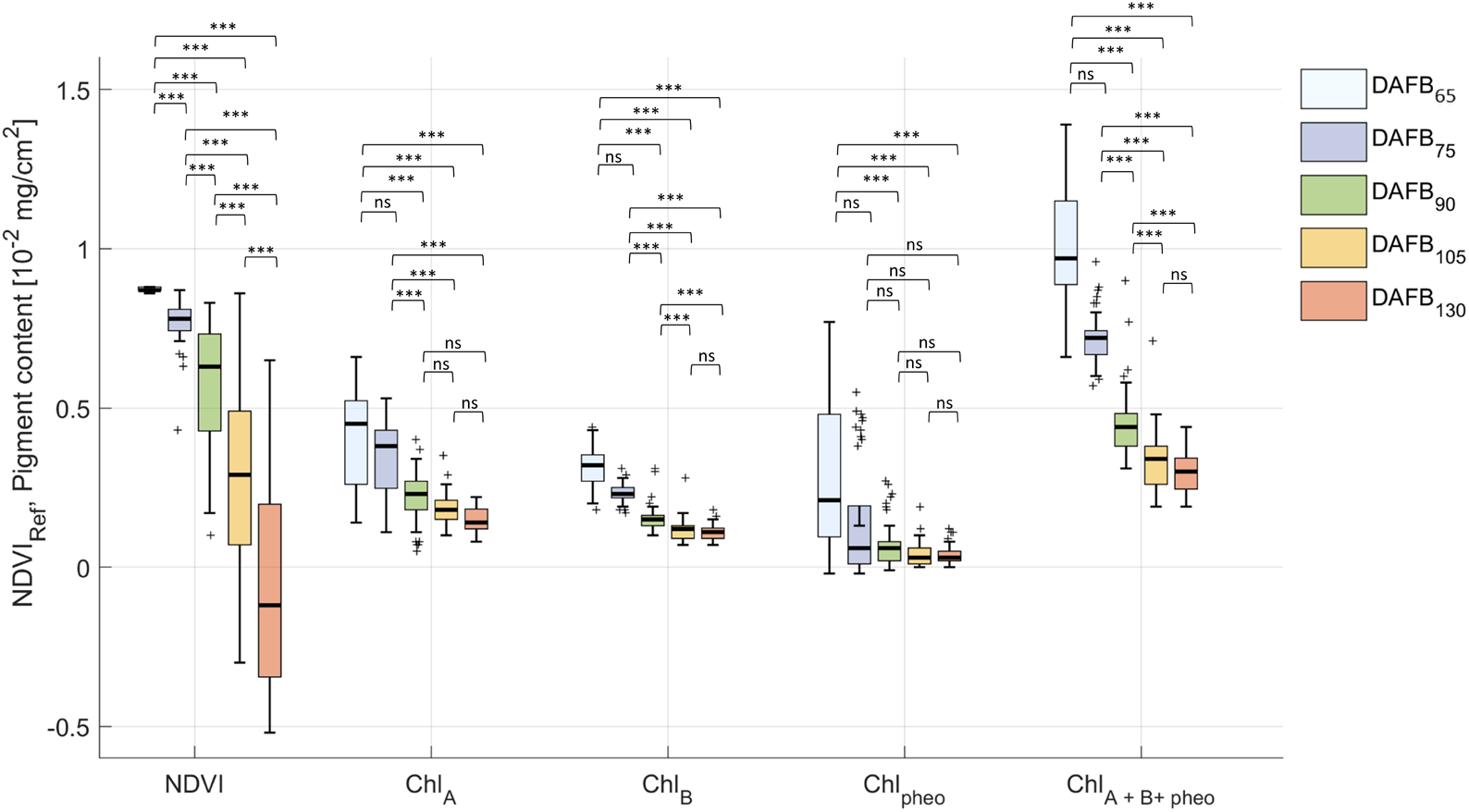
Fruit chlorophyll content and NDVI during fruit development in days after full bloom (DAFB) (n = 45, total = 270). In each box, the centre line represents the mean value, the top and bottom of the box correspond to the 25th and 75th percentiles, and the whiskers represent the 10th and 90th percentiles. Note: ‘***’ indicates a significant difference between classes (p < 0.001) and ‘ns’ indicates non-significant.

The NDVI detected by means of LiDAR (NDVI_LiDAR_) decreased from 0.36 at DAFB_65_ to 0.13 DAFB_130_ (Table 2). A high relation was observed between NDVI_Ref_ and NDVI_LiDAR_ at DAFB_130_ (R^2^ = 0.85, RMSE = 0.02 %), while low correlations were found at DAFB_65_, DAFB_75_, and DAFB_105_. The Chl_A_ mean value ranged from 0.36 to 0.12 10^-2^ mg/cm^2^ over the growth stages. Generally, NDVI_LiDAR_ showed enhanced correlation with the Chl_A_ compared to other chlorophylls over the growing stages. More specifically, a R^2^ of 0.85, 0.95, 0.82, 0.59, and 0.81 at DAFB_65_ DAFB_75_, DAFB_90_, and DAFB_130_, respectively.

**Table 2.**
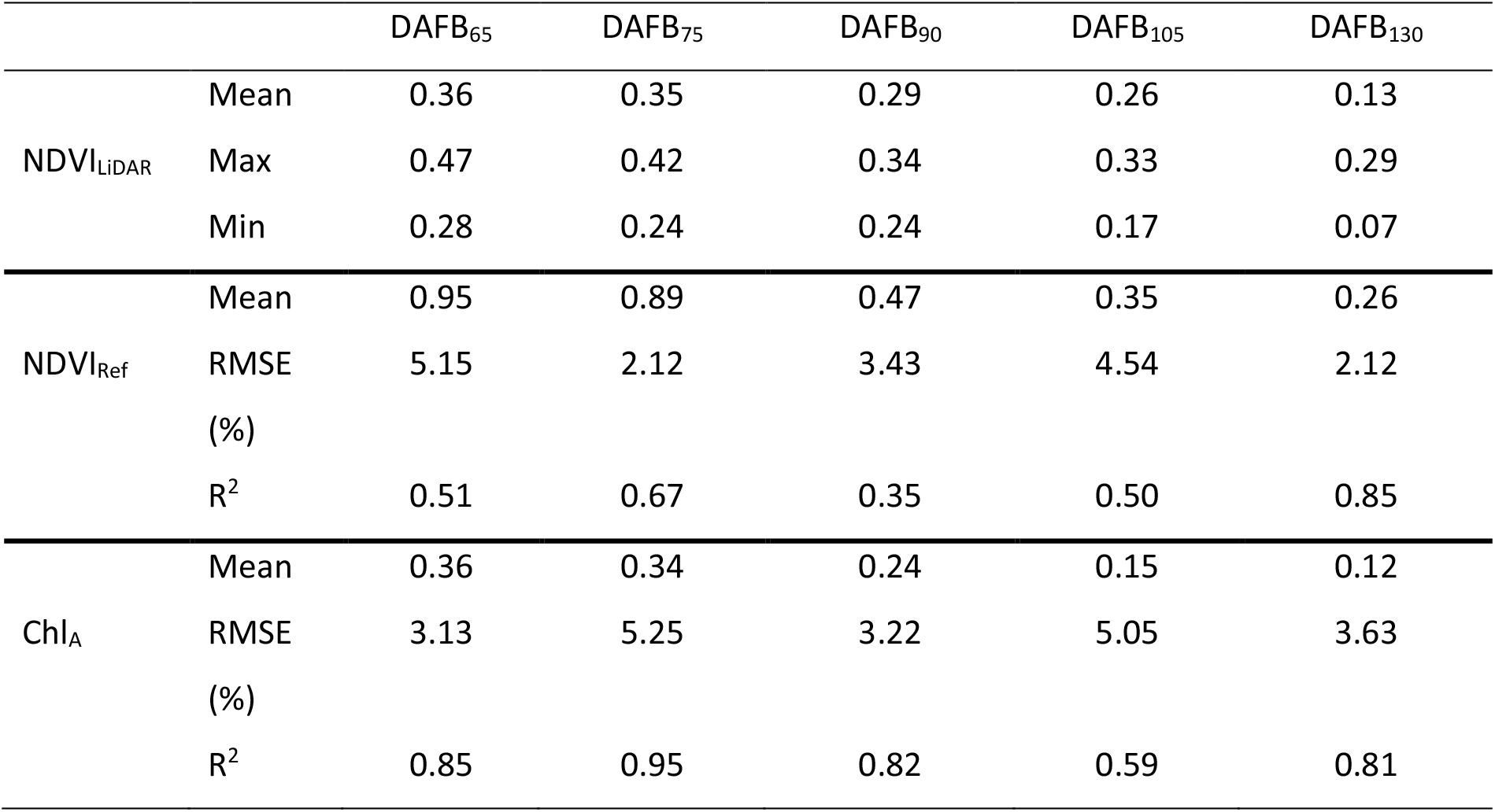
Reference (manual) data of chlorophyll content, NDVI (NDVI_Ref_) and LiDAR derived NDVI estimation (NDVI_LiDAR_) in the laboratory (n =10) with error analysisat five growth stages of apple.

A linear model was used to express the overall relationship between the NDVI_Ref_ and the NDVI_LiDAR_, revealing high R^2^ = 0.86 with RMSE = 0.01 % calculated from the 3D point cloud of apples measured in the laboratory (n = 10) (Figure 7). In parallel, the overal relationship between the Chl_A_ and the NDVI_LiDAR_ was expressed with a logarithmic equation, revealing R^2^ = 0.81 and RMSE = 0.02 considering all growing stages (Figure 7).

**Figure 7:**
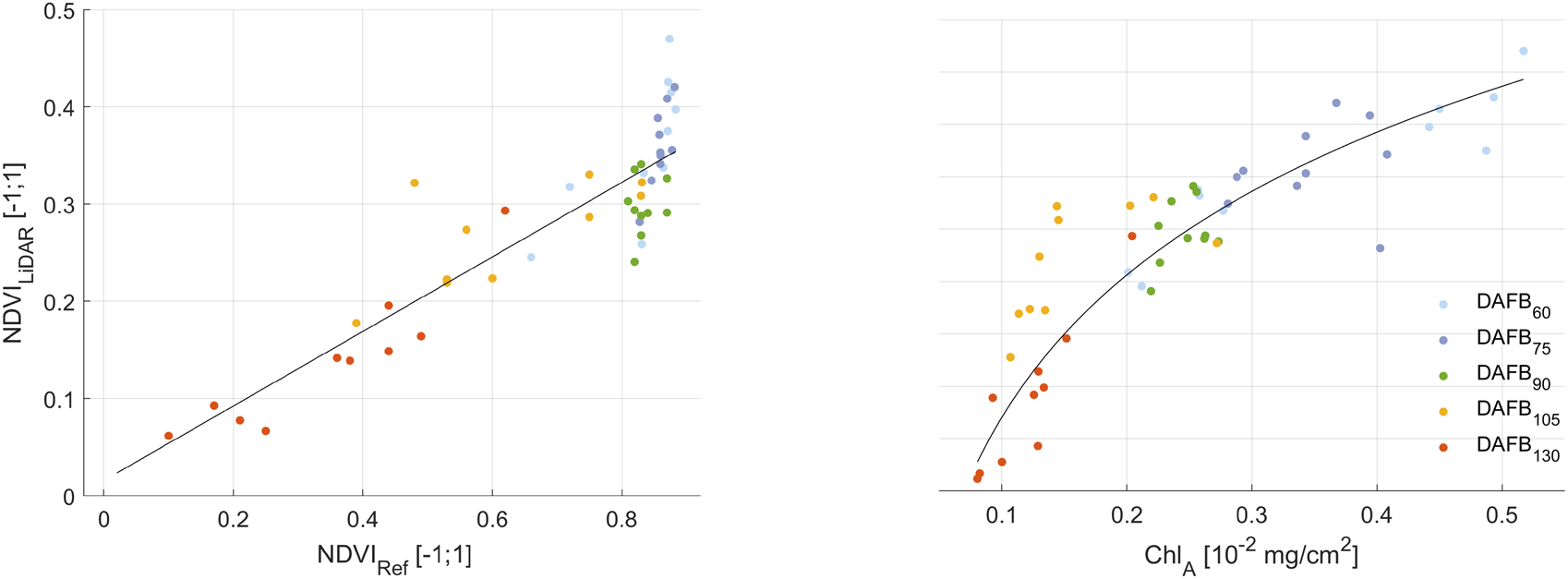
Scatterplots of LiDAR derived NDVI (NDVI_LiDAR_) with (left) manually measured NDVI_Ref_ and, (right) chlorophyll content at five ripening stages of apples measured in the laboratory (n = 10).

In contrast to the laboratory measurements, mean values of NDVI_LiDAR_ measured in the orchard showed higher range (Table 3). However, low and moderate correlations observed over the growth stages with the highest to occure at DAFB_75_ (R^2^ = 0.51, RMSE = 0.02 %) and DAFB_90_ (R^2^ = 0.61, RMSE = 0.02 %).

**Table 3.** LiDAR derived NDVI (NDVI_LiDAR_, n = 12) measured in the orchard with error analysis considering NDVI measured with spectrophotometer in contact to the fruit at the tree (NDVI_Ref_) at five growth stages of apple (total = 60).

| | | $DAFB_{65}$ | $DAFB_{75}$ | $DAFB_{90}$ | $DAFB_{105}$ | $DAFB_{130}$ |
| --- | --- | --- | --- | --- | --- | --- |
| $NDVI_{LiDAR}$ | Mean | 0.95 | 0.89 | 0.47 | 0.35 | 0.26 |
|  | RMSE | 1.54 | 1.58 | 1.54 | 2.87 | 2.26 |
| | $R^2$ | 0.41 | 0.51 | 0.61 | 0.12 | 0.37 |

The NDVI_LiDAR_, measured in the orchard, was evaluated with the manually measured NDVI (NDVI_Ref_) in low and high sections of the tree canopy (Figure 8). The fruit measured in low section below 1.8 m, revealed R^2^ = 0.62 with RMSE = 0.1 % (Figure 8, left), while a R^2^ = 0.71 with RMSE = 0.1 % RMSE was found (Figure 8, right). Furthermore, the Chl_A_ was measured in low and high sections of canopies, resulting in R^2^ = 0.41 with RMSE = 1. 46 and R^2^ = 0.81 with RMSE = 1.18 % at harvest stage (Figure 9), respectively.

**Figure 8:**
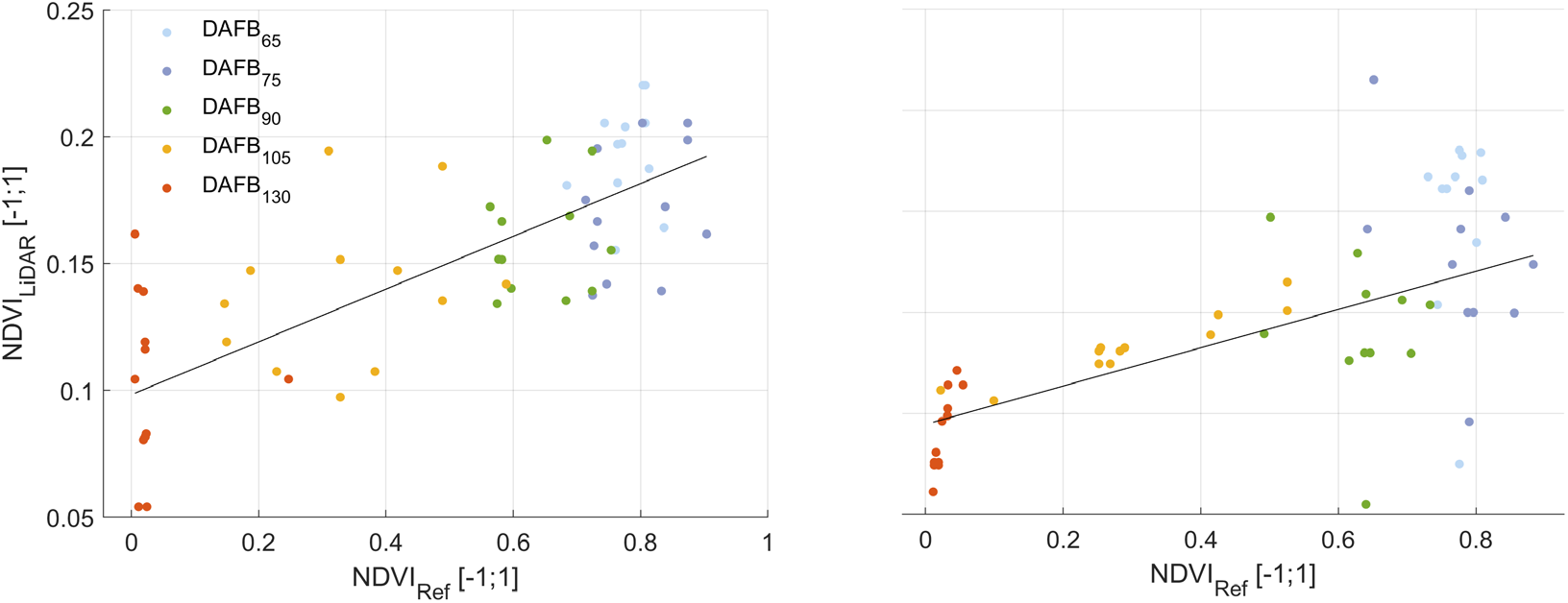
Scatter plots of LiDAR derived NDVI (NDVI_LiDAR_) and manually measured NDVI_Ref_ (n = 12) in (left) low and (right) high areas of the tree measured in the field, at five ripening stages of apples.

**Figure 9.**
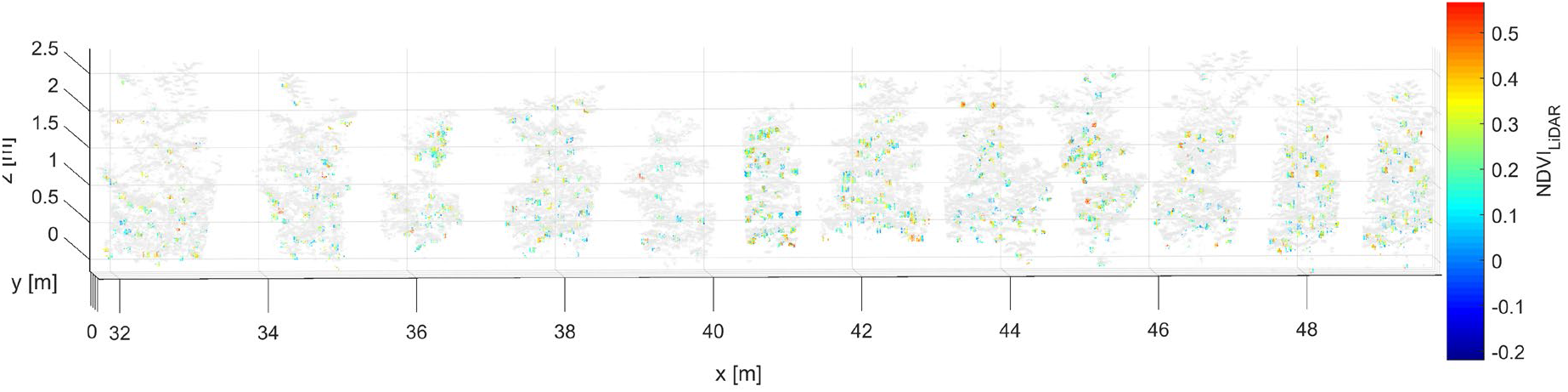
3D point clouds of 12 apple trees with the normalised vegetation index estimated by means of LiDAR (NDVI_LiDAR_) in apples, measured on the last measuring date at harvest.

### Impact of crop load on fruit NDVI

The mean of segmented leaf area by means of LiDAR analysis (LA_LiDAR_) increased from 5.85 m^2^ to 5.98 m^2^ during measuring period in upper section (> 1.8 m) of the tree, while lower tree section (< 1.8 m) developed higher LA_LiDAR_ with 6.17 m^2^ and 6.44 m^2^ between DAFB_65_ and DAFB_130_, respectively (Figure S1). Destructive measurements of LAManual and points per tree (PPT) excluding points of wood were used to build a linear regression model for estimating the LA of each tree from the 3D tree point clouds (Figure S1). including all phenological stages, LA_LiDAR_ showed an R^2^ of 0.87 with 1.32 % RMSE and an R^2^ 0.96 with 0.83 %, in low and high areas of the tree, respectively.

The segmented leaf area and number of fruit were used to estimate the leaf area to fruit ratio (LA_LiDAR_: Fruit_LiDAR_) in low and and high section of tree canopy (Figure 10). The LA_LiDAR_: Fruit_LiDAR_ varied from 0.12 m^2^ fruit^-1^ at DAFB_65_ to 0.17 m^2^ fruit^-1^ at DAFB_130_ in low areas, while a higher range was found in the upper part of canopies reaching 0.31 m^2^ fruit^-1^ at harvest. Clear difference was observed (p < 0.001) among the growth stages of the two canopy sections. The highest percentage difference was found at harvest stage reaching 58.3 %.

**Figure 10.**
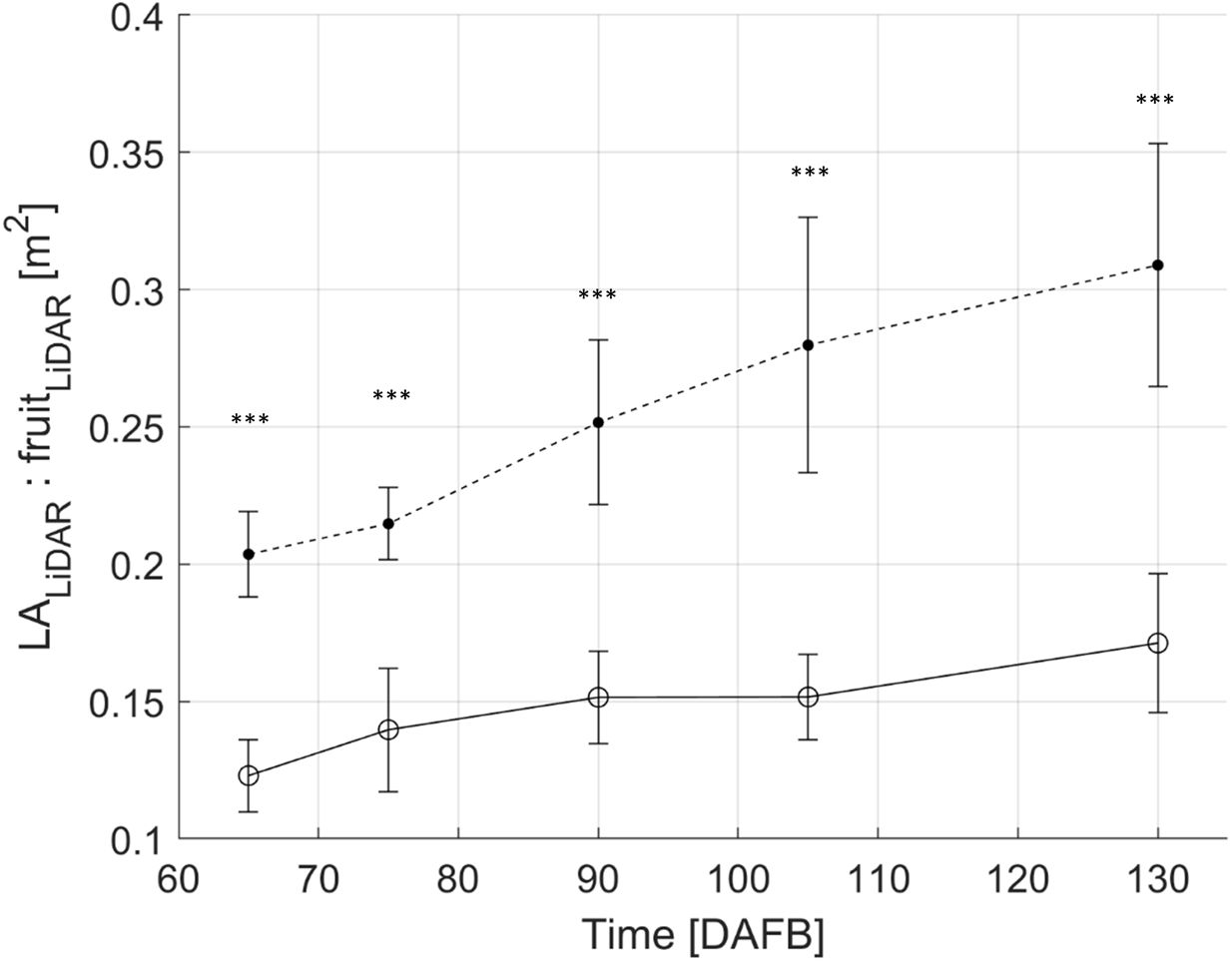
Line graph of temporal development of leaf area to fruit ratio by means of LiDAR (LA_LiDAR_ and Fruit_LiDAR_) in low (blank) and high (filled) section of the canopy (n = 12) over fruit growth stages. Note: ‘***’ indicates a significant difference between classes (p < 0.001).

The individual fruit NDVI_LiDAR_ showed no interaction with LA_LiDAR_: Fruit_LiDAR_ over the growth stages. However, a logarithmic model was able to express the relationship of the two variables, when considering the average values of each measuring date (Figure 11). The LA_LiDAR_: Fruit_LiDAR_ correlated with the NDVI_LiDAR_, revealing R^2^ = 0.71 and RMSE = 2.86 % in the upper canopy section, while a similar R^2^ = 0.74 with RMSE = 2.46 % was observed in the lower section.

**Figure 11.**
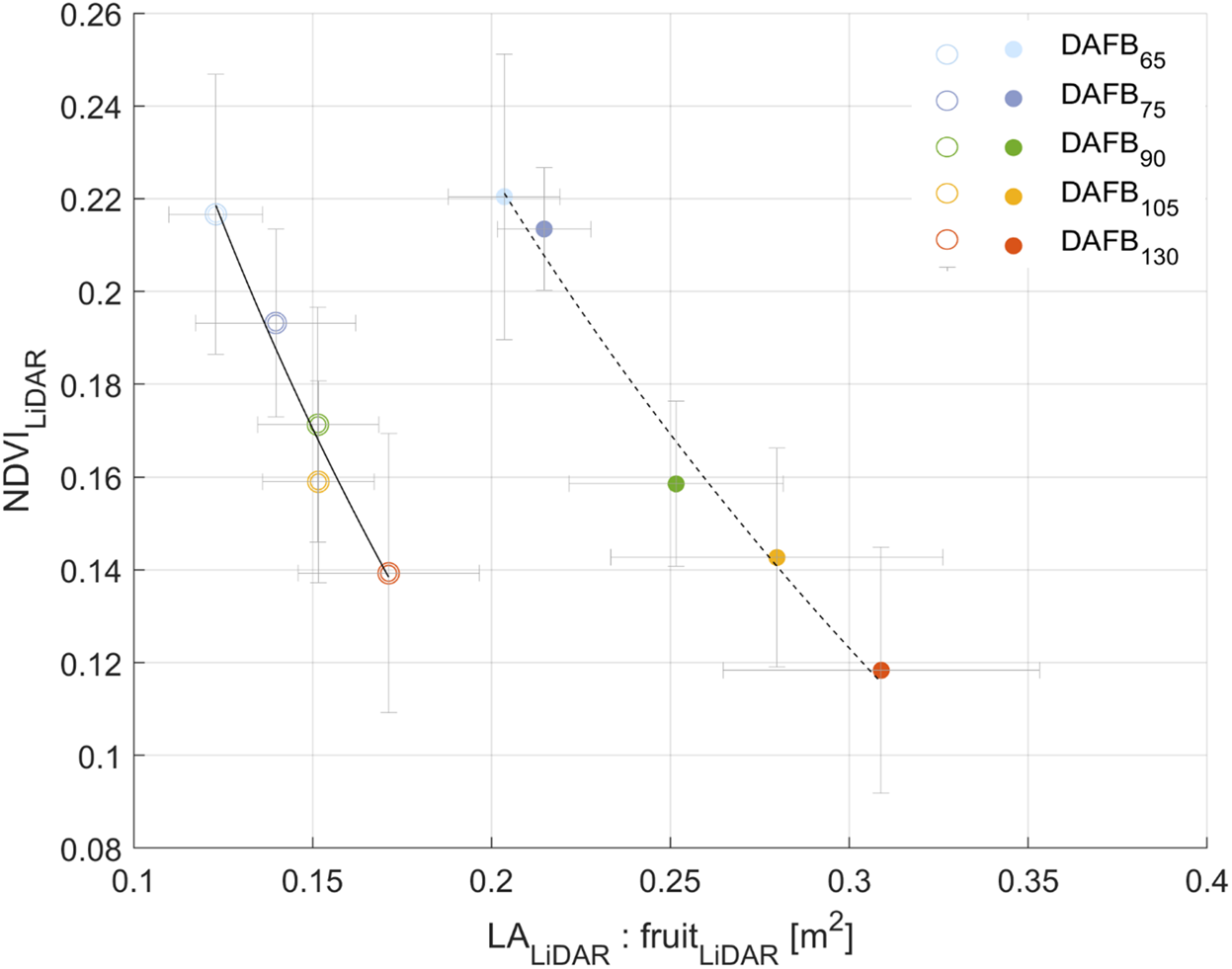
Regression analysis of LiDAR detected leaf area to fruit number (LA_LiDAR_: Fruit_LiDAR_) and NDVI (NDVI_LiDAR_) in low (< 1.8 m, open) and high (> 1.8 m, filled) section of the tree (n =12) over fruit development stages in day after full blum (DAFB).

The manually measured fresh mass correlated with the manually measured diameter (D_Ref_) with an R^2^ of 0.95 and RMSE = 0.86 %, while the diameter derived by means of LiDAR laser scanner (D_LiDAR_) showed a similar R^2^ of 0.94 with RMSE = 0.96 % (Figure S2). A steep decrease of LA_LiDAR_: FM_LiDAR_ found between the DAFB_65_ and DAFB_75_ in low section of the tree (Figure S3). In general, less pronounced values were observed in the upper part, revealing a mean value of 0.22 m^2^ g^-1^ at DAFB_65_ and 0.03 m^2^ g^-1^ at DAFB_130_ (Figure 12).

**Figure 12.**
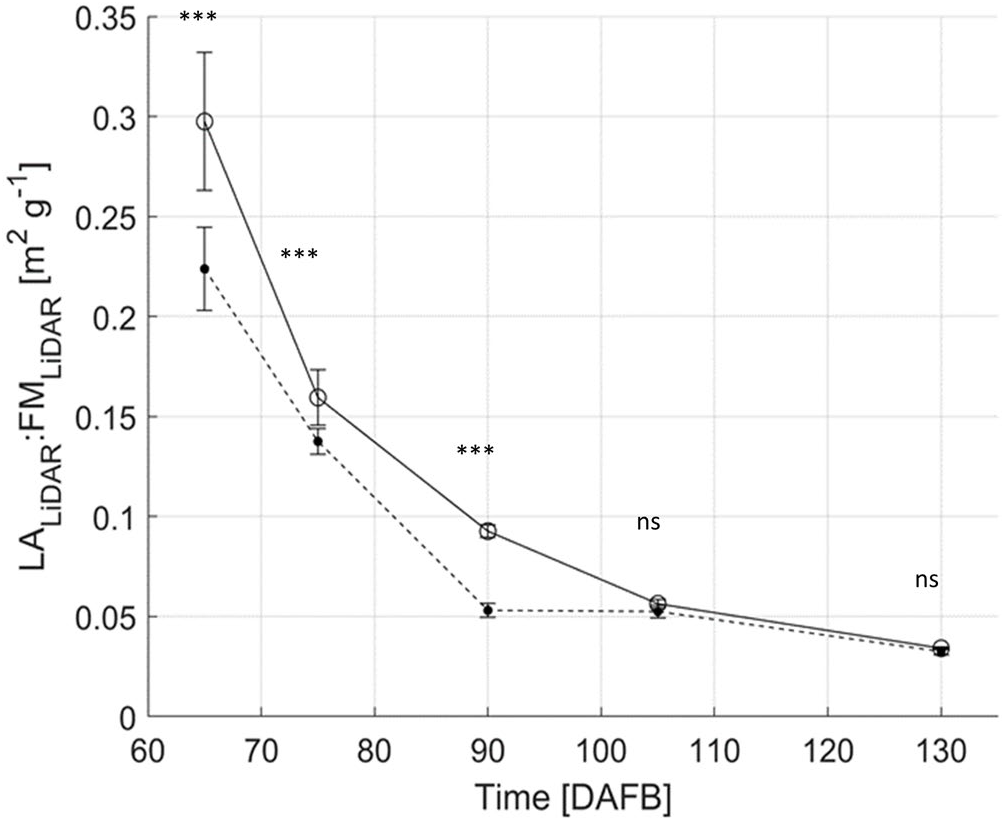
Line graph of temporal development of leaf area to fruit fresh mass ratio by means of LiDAR (LA_LiDAR_:FM_LiDAR_) in low (blank) and high (filled) section of canopy over fruit development stages (n =12). Note: ‘***’ indicates a significant difference between classes (p < 0.001) and ‘ns’ indicates non-significant.

In contrast to the LA_LiDAR_: Fruit_LiDAR_, moderate and high relations were observed between LA_LiDAR_: FM_LiDAR_ and NDVI_LiDAR_ in most growth stages (Table 4). More specifically, R^2^ of 0.51, 0.91, and 0.71 was found at DAFB_75_, DAFB_105_ and DAFB_150_ in high canopy section, respectively. On the other hand, lower parts of the tree revealed enhanced correlations of less pronounced uncertainty 0.83 – 1.16 %, 0.85 – 2.06 %, 0.57 – 1.17 % and 0.83 – 1.15 % at DAFB_65_, DAFB_75_, DAFB_90_ and DAFB_150_, respectively.

**Table 4:**
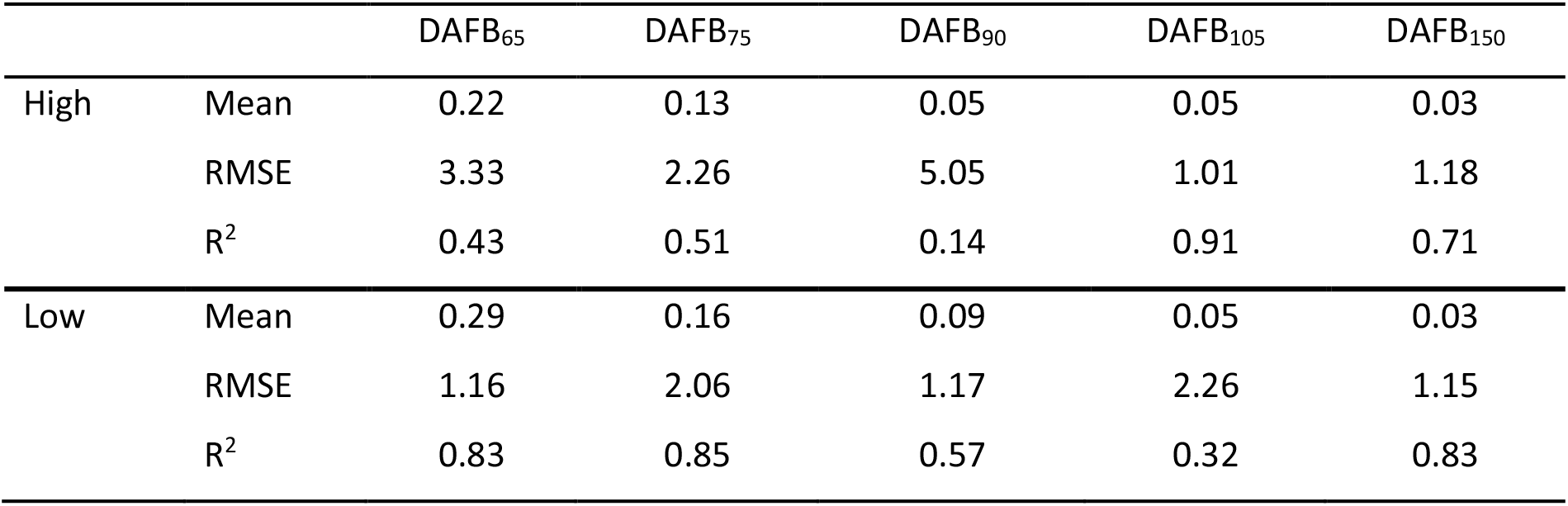
Descriptive statistics of LiDAR derived fruit NDVI (NDVI_LiDAR_) and leaf area to fruit fresh mass ratio (LA_LiDAR_: FM_LiDAR_) (n =12) measured in the field in low (<1.8 m) and high (>1.8 m) sections of the tree at five growth stages of apple.

The NDVI_LiDAR_ correlated with the LA_LiDAR_: FM_LiDAR_ resulting in an R^2^ of 0.95 and 0.66 % RMSE in high areas, while a similar correlation, R^2^ 0.91 with 0.73 % RMSE, was presented in the lower part (Figure S3). Both curves showed a reciprocal trend, that decreased along with chlorophyl degradation.

Additionally to the analysis of fruit chlorophyll based on fruit NDVI_LiDAR_, also the leaf NDVI was measured. Earlier work showed high correlation of LiDAR data and leaf chlorophyll content (Hosoi et al., 2019; Sun et al., 2019). In the present work such findings were confirmed for apple trees measured during the fruit growth period (Figure S4). The leaf NDVI_LiDAR_ was linearly related with the corresponding Chl_A_ content, resulting in R^2^ = 0.61 and RMSE = 1.32 % and R^2^ = 0.62 with RMSE = 1.12 % in low and high section of canopies, respectively.

## Conclusions

LiDAR 3D point clouds acquired in the dark room and field conditions provided geometric information of apple trees to count and size fruit and leaves. It was found that 660 and 905 nm wavelength of the LiDAR enables to produce the NDVI_LiDAR_ point cloud of apples and leaves. The NDVI_LiDAR_ curves derived in the laboratory and field followed chlorophyll content degradation of apples over the ripening stages. NDVI_LiDAR_ resulted in high coefficient of determination with Chl_A_ (R^2^ = 0.81) and NDVI_Ref_ (R^2^ = 0.86) when measured in dark room. Less pronounced relationship of NDVI_LiDAR_ were observed with NVDIRef in the field, in apples located in low (R^2^ = 0.62) and high (R^2^ = 0.72) sections of canopies. The leaf area to fruit ratio derived by means of LiDAR interacted with the NDVI_LiDAR_ of apples, revealing an R^2^ = 0.74 in the upper section of the tree. The regression curve of LA_LiDAR_: FM_LiDAR_ and NDVI_LiDAR_ also exhibited enhanced correlations of R^2^ = 0.95 and R^2^ = 0.91 in high and low sections of the tree, respectively. Overall, this study shows the applicability of LiDAR backscattered intensity to analyse the NDVI_LiDAR_ and estimate the chlorophyll contents of fruit and leaves. Extracting the crop load, shape, and chlorophyll content parameters can support decision making of apple harvesting robotic platforms and allowing further physiological studies of fruit development during changing climate conditions.

## Acknowledgements

We thank Gabi Wegner and Corinna Rolleczek for manual rating and chemical lab analysis of fruit and acknowledge the technical support of Christian Regen in the setup of sensors on the conveyors and coding of data acquisition routine.

The project SPECTROFOOD is part of the ERA-NET Cofund ICT-AGRI-FOOD, with funding provided by national sources of BMEL and co-funding by the European Union’s Horizon 2020 research and innovation program, Grant Agreement number 2820ERA16H. This publication was supported by the Open Access Publication Fund of the Leibniz Association.

## Supplement

**Figure S1.**
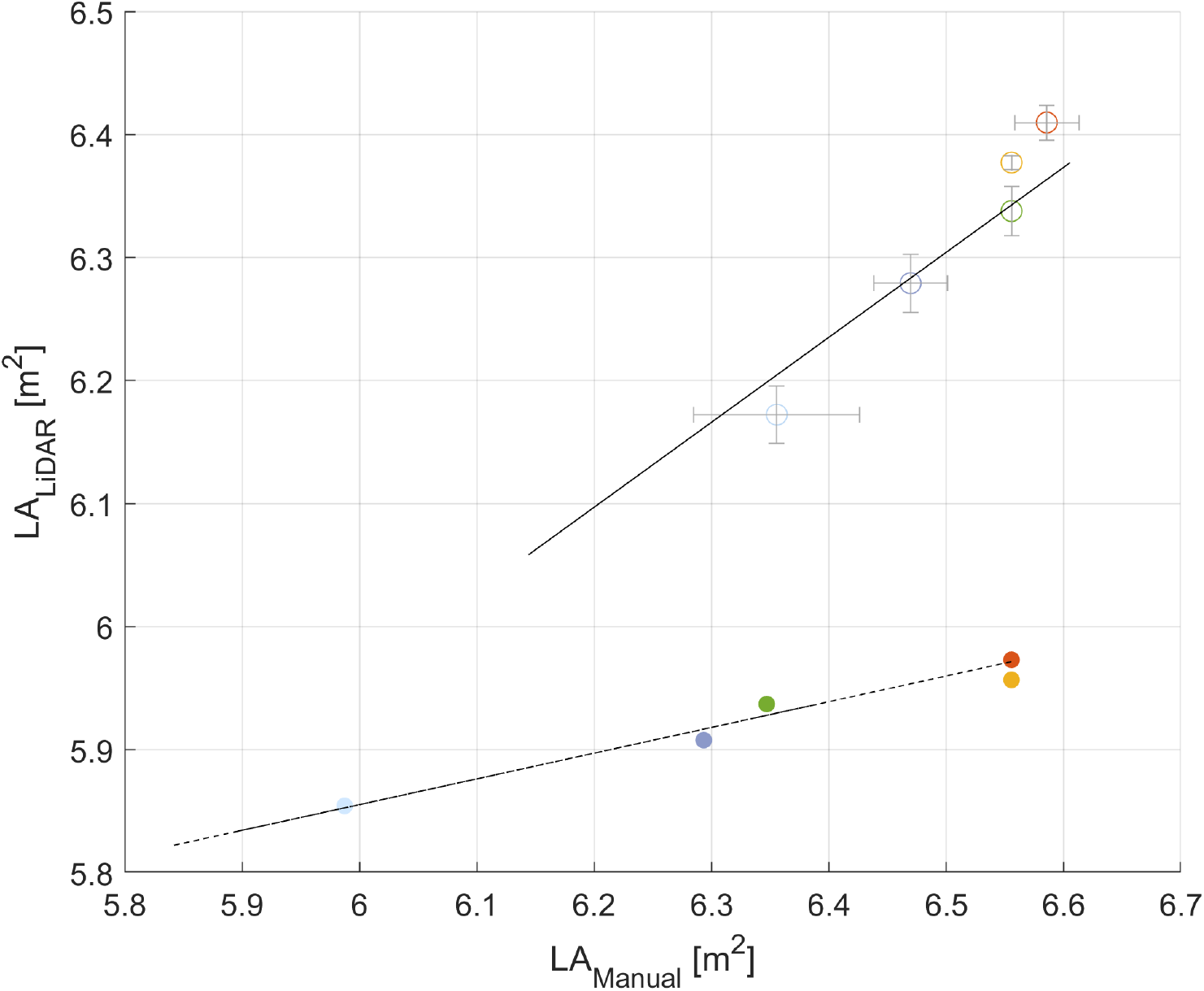
Scatterplot of manually measured leaf area (LAManual) with LiDAR detected leaf area (LA_LiDAR_) in low (blank) and high (filled) section of tree canopy over fruit development stages (n = 36).

**Figure S2.**
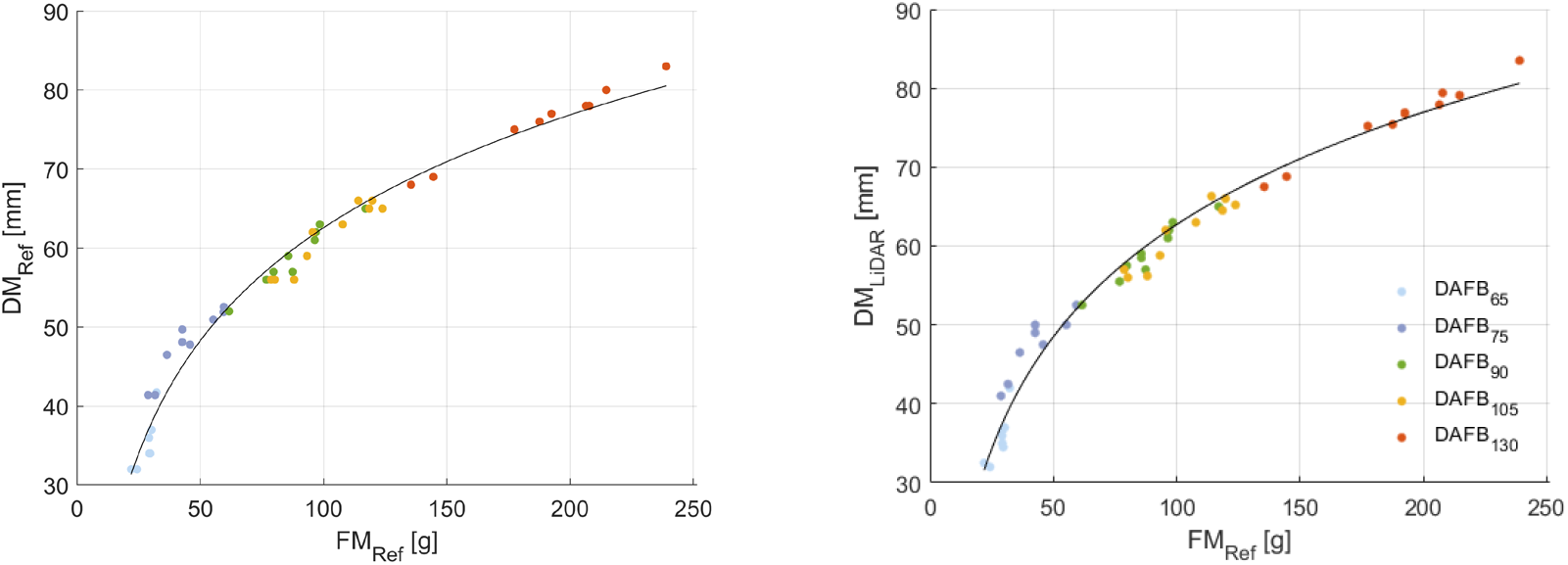
Scatter plots of manually measured fresh mass with (left) manually measured diameter and (right) LiDAR derived diameter (DM_LiDAR_) (n = 10, total = 60) from apples measured in the laboratory at five growth stages.

**Figure S3.**
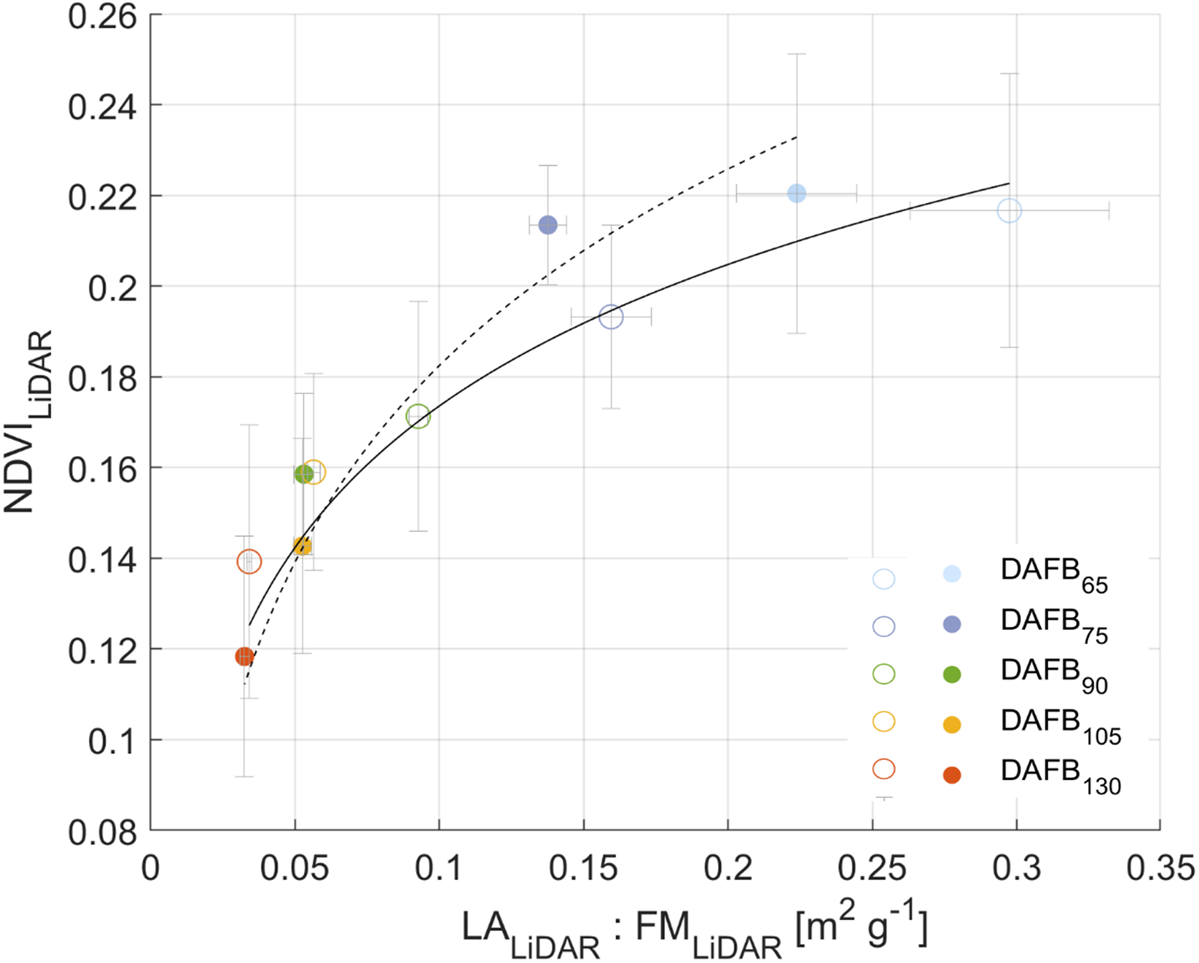
Scatterplot of LiDAR detected leaf area to fruit ratio (LA_LiDAR_: FM_LiDAR_) and NDVI (NDVI_LiDAR_) in low (blank) and high (filled) section of the tree (n =12) over fruit growth stages.

**Figure S4.**
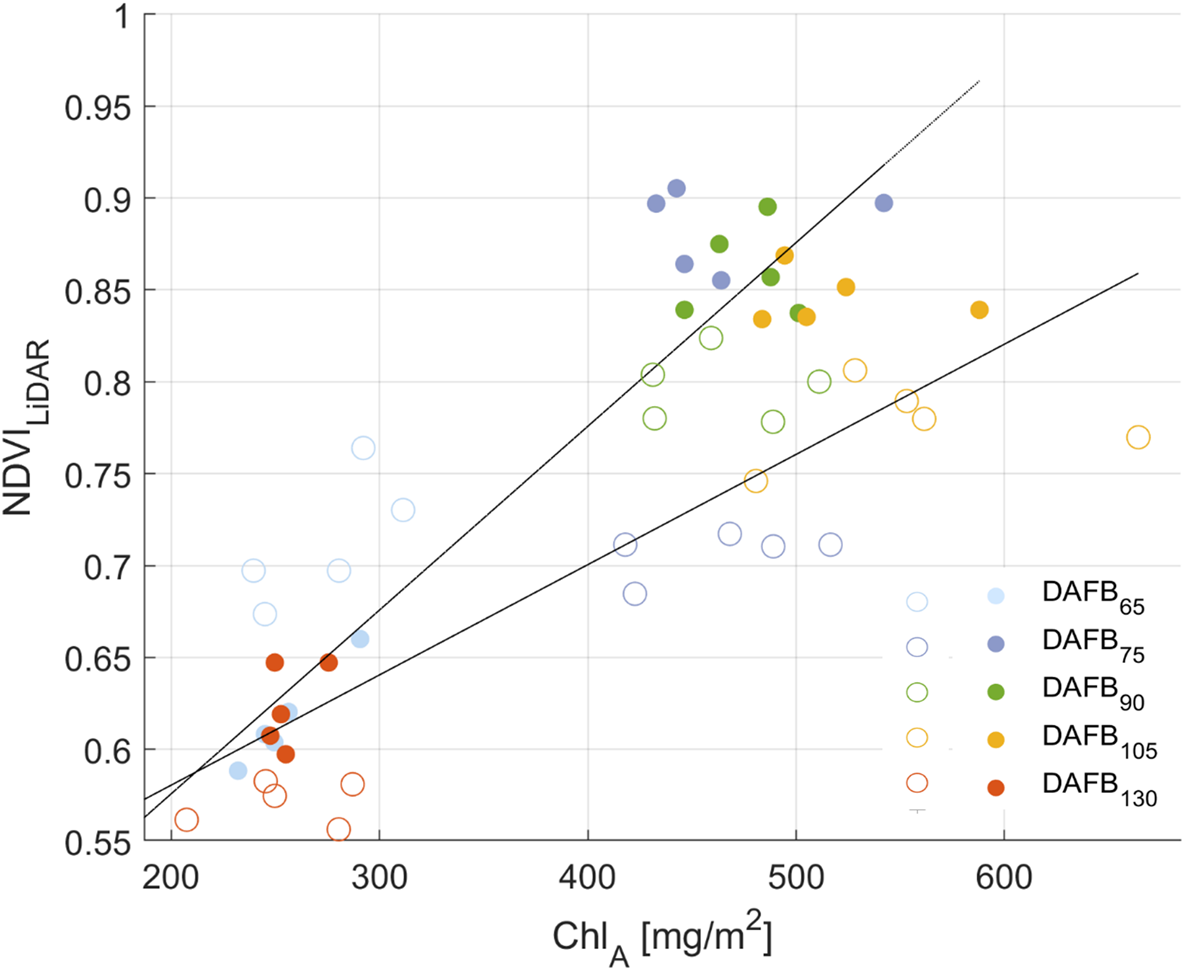
Scatterplot of manually measured chlorophyll of leaf area (Chl_A_) with LiDAR detected leaf area NDVI (NDVI_LiDAR_) in low (open) and high (closed) section of the tree over fruit development stages.

